# Comparative analysis of mammalian adaptive immune loci revealed spectacular divergence and common genetic patterns

**DOI:** 10.1101/2025.04.01.646651

**Authors:** Mariia Pospelova, Katalin Voss, Anton Zamyatin, Corey T. Watson, Klaus-Peter Koepfli, Anton Bankevich, Matt Pennell, Yana Safonova

## Abstract

Adaptive immune responses are mediated by the production of adaptive immune receptors, antibodies and T-cell receptors, that bind antigens, thus causing their neutralization. Unlike other proteins, adaptive immune receptors are not fully encoded in the germline genome and result from a complex of somatic processes collectively called V(D)J recombination affecting germline immunoglobulin (IG) and T-cell receptor (TR) loci consisting of template genes. While various existing studies report extreme diversity of antibodies and T-cell receptors, little is known about the diversity of germline IG and TR loci. To overcome this gap, the first comparative analysis of full-length sequences of IG/TR loci across 44 mammalian species from 13 taxonomic orders was performed. First, germline genes counts were shown to correlate in IGH/IGL and TRA/TRB and anticorrelate in IGK/IGL, possibly indicating co-evolution between corresponding chains. Second, structures of IG/TR loci were analyzed, and it was shown that IG/TR loci formed by long arrays of high multiplicity repeats are more common for species that have experienced population bottlenecks. Finally, haplotypes of IG/TR loci with little or no sequence similarity within a species were found, suggesting that they may have a limited potential for homologous recombination. These results demonstrate that IG/TR loci are rapidly evolving genomic regions whose structural variation is shaped by the population history of the species and open new perspectives for immunogenomics studies.

## Introduction

Adaptive immune responses are mediated by the production of antibodies (Abs) and T-cell receptors (TCRs) produced by B- and T-cells, respectively, that bind to bacteria, viruses, or aberrant cells (collectively called antigens) and cause their neutralization or destruction. The collection of expressed Abs (TCRs) in an individual is known as the Ab (TCR) repertoire. The generation of diverse Abs and TCRs relies on a process called V(D)J recombination that somatically rearranges the DNA in individual B- and T-cells. In B-cells, this process affects germline *immunoglobulin* (*IG*) loci, in which individual IG genes including variable (V), diversity (D), and joining (J) genes are selected and generate a VDJ sequence corresponding to *the variable region* for either the heavy chain (HC) or the light chain (LC) of a given antibody (Tonegawa, 1983). Most mammalian genomes contain one IG heavy chain locus (IGH) and two immunoglobulin light chain loci: kappa (IGK) and lambda (IGL). Light chain loci do not contain D genes and generate VJ recombinations. In T-cells, the V(D)J recombination process affects germline *T-cell receptor* (*TR*) loci. Most mammalian species contain four types of TR loci: TR alpha (TRA), TR beta (TRB), TR gamma (TRG), and TR delta (TRD) loci encoding different types of TR chains. TRA and TRD loci contain V and J genes only, and TRB and TRG contain V, D, and J genes (Janeway et al., 2001).

While existing studies of adaptive immune repertoires note their extraordinary diversity resulting from V(D)J recombination (Alberts et al., 2022; Schroeder, 2006; Briney et al., 2019; Safonova et al., 2022), little is known about the structure of germline IG/TR loci encoding them. The main reason for this gap is high repetitiveness of IG/TR loci which makes them particularly difficult to reconstruct from short-read sequencing reads and often leads to fragmented assemblies (Watson et al., 2013; Rodriguez et al., 2020; Lin et al., 2024; Zhu et al., 2024). Since incomplete assemblies prevent a comprehensive analysis of germline sequences, comparative immunogenomics studies have instead focused on individual germline IG/TR genes (Ramesh et al., 2017; Luo et al., 2019; Kaduk et al., 2022; Sirupurapu et al., 2022; Pennell et al., 2023; Ma et al., 2024), which can still be identified in fragmented assemblies. These studies reported the remarkable sequence diversity of germline IG/TR genes across species, thus showing that the diversity of Ab/TCR repertoires is built upon it and is further enhanced by the somatic V(D)J recombination process.

Recent advances in long-read sequencing technologies and genome assembly methods have enabled nearly complete reconstruction of haplotype-resolved sequences of IG/TR loci, however, genetic characteristics and architecture of germline IG/TR loci remains poorly studied. Most existing comparative immunogenomics studies either focused on direct comparisons with human IG/TR loci (e.g., human and mouse IGH: Sepulveda et al., 2005; dog and human IGL: Martin et al., 2018) or studied a specific clade. For example, Rodriguez et al., 2023 analyzed germline human IGH loci and described novel structural variations across 150 individuals. Sampson and Miller, 2023 investigated the impact of mobile elements in marsupial IG loci. Yoo et al., 2024 characterized germline IG and TR loci of four great ape species and revealed that structural variations harbor specific-specific IG and TR genes. Pursell et al., 2024 analyzed bat IG loci and discovered an IGH locus duplication. Despite these advances, no study has conducted a broad comparative analysis of IG/TR loci across diverse clades. A major challenge lies in the rapid evolutionary divergence of these loci, which results in little to no sequence similarity across species. The inability to compute alignment and identify orthologous genes makes these regions difficult to analyze using comparative genomics tools such as IGV (Robinson et al., 2011) or SyRi (Goel et al., 2019) and raises a need in new approaches handling rapidly evolving regions.

In this paper, we attempted to close this gap and performed the first comparative study of full-length haplotype-resolved sequences of germline IG/TR loci across 44 therian mammalian species spanning 13 taxonomic orders. We combined the high-quality data generated from large-scale genome sequencing projects and developed a set of metrics describing germline gene diversity, haplotype divergence, and genomic architecture of IG/TR loci. We revealed associations between these metrics and found connections with species phenotypes.

## Results

### Predicting adaptive immune loci in mammalian assemblies

High-quality diploid genome assemblies of 44 therian mammalian species (42 placental and 2 marsupial mammals) representing 13 taxonomic orders were collected from the Vertebrate Genomes Project (Rhie et al., 2021; www.genomeark.org); the California Conservation Genomics Project (Shaffer et al., 2022; www.ccgproject.org), and the Bovine Pangenome Consortium (Smith et al., 2023; bovinepangenome.github.io) (**Fig. 1A**). For 3 out of 44 species (the cattle, the European badger, the black rhinoceros), parental information via trios was available and the genomes were haplotype phased; for one species (the brown bear), haplotypes were not separated; and for remaining 40 species, genomes were not phased, but both primary and alternate assemblies were available (**Table 1**). Each assembly except for that of the brown bear was marked with a haplotype label (primary/alternate; haplotype1/haplotype2; or maternal/paternal). To predict germline genes and determine the boundaries of IG/TR loci, the IgDetective tool (Sirupurapu et al., 2022) was modified to handle all types of adaptive immune loci and applied to the selected genomes (see “Modifying IgDetective and filtering broken IG/TR loci” in Methods). IG/TR genes were detected in both haplotype assemblies of each species, and, as a result, IGH, IGK, IGL, TRA, and TRB loci were predicted. TRG loci were predicted only for 9 species and were excluded from the analysis. TRA and TRD loci overlap and were not distinguished. The analysis was focused on V genes because they are easier to compare within and across species than D genes and are more numerous than J genes. We classified a V gene as productive if it was in-frame and did not contain internal stop codons. V genes that were detected by IgDetective but did not meet these criteria were classified as pseudogenes.

**Figure 1.**
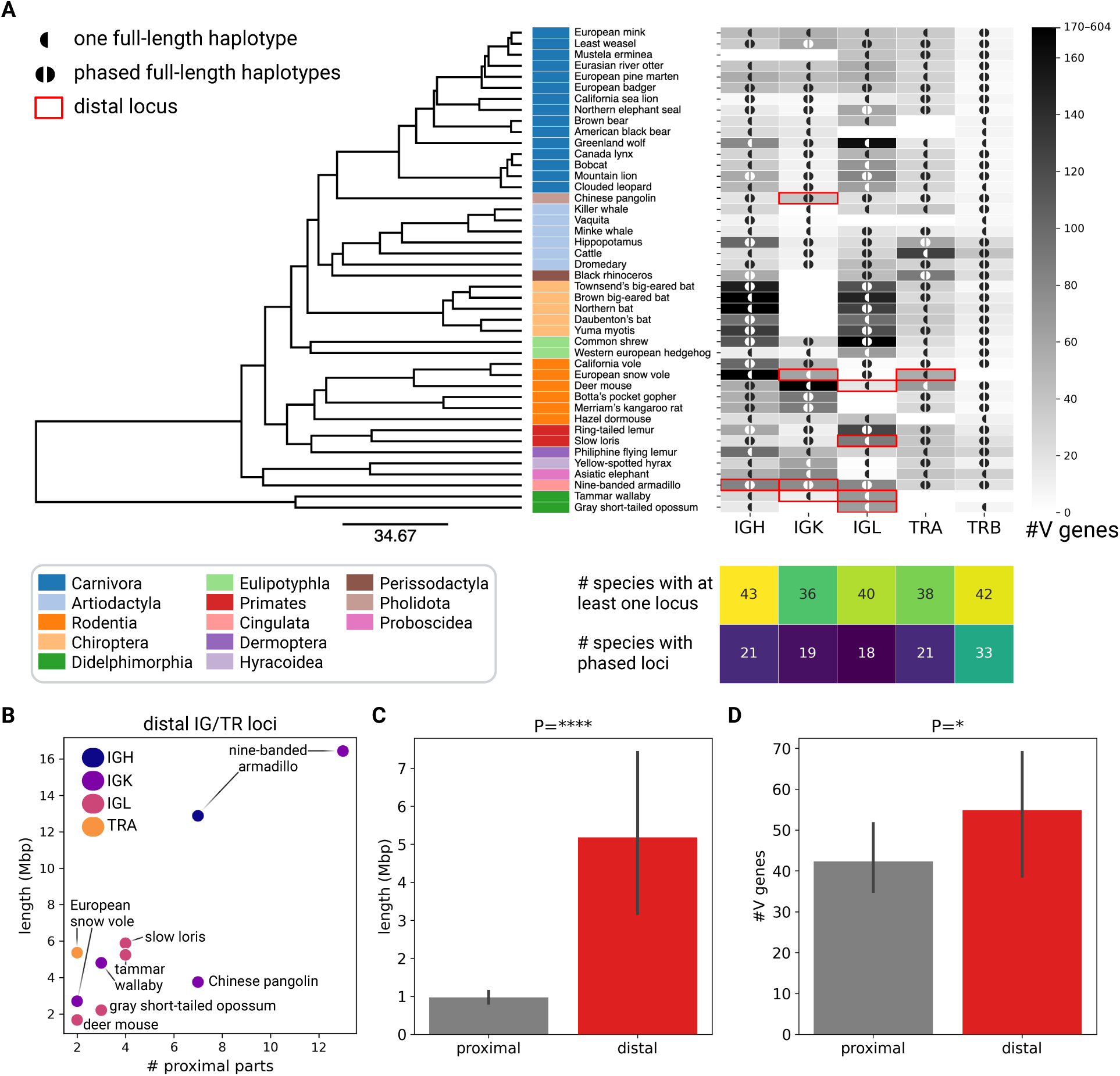
Overview of IG/TR loci collected across 44 mammalian species. **(A)** The phylogenetic tree of mammalian species selected for the analysis. The tree was constructed using consensus topologies derived from 100 trees obtained from VertLife.org (Upham et al., 2019). The final consensus tree was generated with TreeAnnotator v1.10.4 (Drummond et al., 2007) using mean node heights. Taxonomic orders are color coded and shown next to the species names. Counts of productive V genes in IGH, IGK, IGL, TRA, and TRB loci are shown on the heatmap on the right. Distal IG/TR loci are shown as red rectangles. A locus is marked with ◖/◖◗ if it has one/two intact haplotype(s) assembled. Counts of such sequences are summarized in the heatmap below. **(B)** Number of proximal parts vs the total locus length across 10 distal IG/TR loci. **(C)** Lengths of proximal and distal IG/TR loci. Here and further error-bars represent 95% confidence intervals; P-values are denoted as follows: ns: P≥0.05; *<0.05, **<0.01, ***<0.001, ****<0.0001; and P-values are computed using the Kruskal-Wallis test unless specified otherwise. **(D)** Counts of productive V genes in proximal and distal IG/TR loci.

**Table 1.** Species summary with accession numbers for assemblies.

| Order | Sample ID | Latin Name | Common Name | Haplotype1 | Haplotype 2 |
| --- | --- | --- | --- | --- | --- |
| Artiodactyla | mBalAcu1 | *Balaenoptera acutorostrata* | Minke whale | GCA_949987535.1 | GCA_950005055.1 |
| Artiodactyla | mBosTau | *Bos taurus* | Cattle (Piedmontese) | 10.5281/zenodo.5906579 |  |
| Artiodactyla | mOrcOrc1 | *Orcinus orca* | Killer whale | GCA_937001465.1 | GCA_937001455.1 |
| Artiodactyla | mPhoSin1 | *Phocoena sinus* | Vaquita | GCA_008692025.1 | GCA_008692045.1 |
| Artiodactyla | mCamDro1 | *Camelus dromedarius* | Dromedary | genomeark.org/genomeark-all/Camelus_dromedarius.html |  |
| Artiodactyla | mHipAmp2 | *Hippopotamus amphibius kiboko* | Hippopotamus | GCA_030028035.1 | GCF_030028045.1 |
| Carnivora | mCanLor1 | *Canis lupus orion* | Greenland wolf | GCA_905319855.2 | GCA_905319845.1 |
| Carnivora | mLutLut1 | *Lutra lutra* | Eurasian river otter | GCA_902655055.2 | GCA_902653095.1 |
| Carnivora | mLynCan4 | *Lynx canadensis* | Canada lynx | GCA_007474595.2 | GCA_007474575.2 |
| Carnivora | mMelMel3 | *Meles meles* | European badger | GCA_922990625.1 | GCA_922984935.2 |
| Carnivora | mMusErm1 | *Mustela erminea* | Stoat | GCA_009829155.1 | GCA_009829165.1 |
| Carnivora | mNeoNeb1 | *Neofelis nebulosa* | Clouded leopard | GCA_028018385.1 | GCA_028018655.1 |
| Carnivora | mZalCal1 | *Zalophus californianus* | California sea lion | GCA_009762305.2 | GCA_009762295.2 |
| Carnivora | mMarMar1 | *Martes martes* | European pine marten | genomeark.org/vgp-all/Martes_martes.html |  |
| Carnivora | mMusLut2 | *Mustela lutreola* | European mink | genomeark.org/genomeark-all/Mustela_lutreola.html |  |
| Carnivora | mPumCon1.1 | *Puma concolor* | Mountain lion | GCA_028749985.3 | GCA_028749965.3 |
| Carnivora | mMusNiv1 | *Mustela nivalis* | Least weasel | genomeark.org/genomeark-all/Mustela_nivalis.html |  |
| Carnivora | mUrsAme1.0 | *Ursus americanus* | American black bear | GCA_024610735.1 | GCA_024610745.1 |
| Carnivora | UrsArc2.0 | *Ursus arctos* | Brown bear | GCF_023065955.2 | NA |
| Carnivora | mLynRuf1 | *Lynx rufus* | Bobcat | GCF_022079265.1 | GCA_022079275.1 |
| Carnivora | mMirAng1 | *Mirounga angustirostris* | Northern elephant seal | GCA_029215605.1 | GCA_029215635.1 |
| Chiroptera | mMyoDau2 | *Myotis daubentonii* | Daubenton's bat | genomeark.org/vgp-all/Myotis_daubentonii.html |  |
| Chiroptera | mPleAur1 | *Plecotus auritus* | Brown big-eared bat | genomeark.org/vgp-all/Plecotus_auritus.html |  |
| Chiroptera | mEptNil1 | *Eptesicus nilssonii* | Northern bat | genomeark.org/vgp-all/Eptesicus_nilssonii.html |  |
| Chiroptera | mCorTow1.0 | *Corynorhinus townsendii* | Townsend's big-eared bat | GCA_026230055.1 | GCA_026230045.1 |
| Chiroptera | mMyoYum1.0 | *Myotis yumanensis* | Yuma myotis | GCA_028538775.1 | GCA_028536395.1 |
| Cingulata | mDasNov1 | *Dasypus novemcinctus* | Nine-banded armadillo | genomeark.org/genomeark-all/Dasypus_novemcinctus.html |  |
| Dermoptera | mCynVol1 | *Cynocephalus volans* | Philippine flying lemur | GCA_027409185.1 | GCA_027409165.1 |
| Didelphimorphia | mMacEug1 | *Macropus eugenii* | Tammar wallaby | GCA_028372415.1 | GCA_028389975.1 |
| Didelphimorphia | mMonDom1 | *Monodelphis domestica* | Gray short-tailed opossum | GCA_027887165.1 | GCA_027917375.1 |
| Eulipotyphla | mSorAra2 | *Sorex araneus* | Common shrew | GCA_027595985.1 | GCA_027595995.1 |
| Eulipotyphla | mEriEur2 | *Erinaceus europaeus* | Western European hedgehog | GCA_950295315.1 | GCA_950295305.1 |
| Hyracoidea | mHetBru1 | *Heterohyrax brucei* | Yellow-spotted hyrax | GCA_028571685.1 | GCA_028571665.1 |
| Perissodactyla | mDicBic1 | *Diceros bicornis* | Black rhinoceros | GCA_020826845.1 | GCA_020826835.1 |
| Pholidota | mManPen7 | *Manis pentadactyla* | Chinese pangolin | GCA_030020395.1 | GCA_030020945.1 |
| Primates | mLemCat1 | *Lemur catta* | Ring-tailed lemur | GCA_020740605.1 | GCA_020740595.1 |
| Primates | mNycCou1 | *Nycticebus coucang* | Slow loris | GCA_027406575.1 | GCA_027406535.1 |
| Proboscidea | mEleMax1 | *Elephas maximus* | Asiatic elephant | GCA_024166365.1 | GCA_024166345.1 |
| Rodentia | mChiNiv1 | *Chionomys nivalis* | European snow vole | GCA_950005125.1 | GCA_950005115.1 |
| Rodentia | mMusAve1 | *Muscardinus avellanarius* | Hazel dormouse | genomeark.org/vgp-all/Muscardinus_avellanarius.html |  |
| Rodentia | mPerMan1.0 | *Peromyscus maniculatus* | Deer mouse | GCA_026167925.1 | GCA_026167925.1 |
| Rodentia | mThoBot1.0 | *Thomomys bottae* | Botta's pocket gopher | GCA_024803745.1 | GCA_024803775.1 |
| Rodentia | mMicCal1.0 | *Microtus californicus* | California vole | GCA_028537955.1 | GCA_028537755.1 |
| Rodentia | mDipMer1.0 | *Dipodomys merriami* | Merriam's kangaroo Rat | GCA_024711535.1 | GCA_024711525.1 |

### Adaptive immune loci can be distal

A *proximal* adaptive immune locus was defined as a sequence spanning V, D, and J genes, where adjacent genes are separated by no more than 300 kbp. Proximal loci with less than three productive V genes were classified as *orphans* and excluded from the analysis (Table S1). If germline genes of the same locus were still found in more than one contig in the assembly, the corresponding locus was considered broken during genome assembly. Several exceptions were made from this rule: 10 loci had several proximal regions located on the same contigs and separated by at least 300 kbp. For these cases, the locus boundaries were redefined based on the positions of the leftmost gene in the leftmost proximal locus and the rightmost gene in the rightmost proximal locus, and the locus was reclassified as *distal*. Previously identified distal IGK loci were reported in humans and gorillas, and distal IGL loci were found in bonobo (Lefranc, 2001; Engelbrecht et al., 2024; Yoo et al., 2024). **Fig. 1A** shows that distal loci emerge across different branches of the mammal evolutionary tree. Fig. S1 showing dot plots of 10 distal IG/TR loci illustrates that long-range inversions likely played a role in the formation of five of them (the Chinese pangolin IGK, the European snow vole TRA, the nine-banded armadillo IGK, the slow loris IGL, the tammar wallaby IGK), and the mechanisms underlying the formation other five distal loci (the deer mouse IGL, the European snow vole IGK, the gray short-tailed opossum IGL, the nine-banded armadillo IGH, the tammar wallaby IGL) are less unclear. The number of proximal regions per distal locus varies from 2 to 13 (**Fig. 1B**), and the lengths of non-coding regions between them vary from 300 to 3677 kbp (Fig. S1). On average, distal adaptive immune loci are longer than the proximal ones (Kruskal-Wallis test, P=6.82×10^−7^, **Fig. 1C**) and encode more V genes (Kruskal-Wallis test, P=0.02, **Fig. 1D**). It is unclear whether all proximal parts of a distal IG/TR locus can participate in V(D)J recombination, however, the fact that they were detected across different taxonomic orders suggests that it is a relatively common phenomena that might contribute to expansion of IG/TR loci. Population-wide genome sequencing data and data from expressed adaptive immune repertoires will help to better understand frequency and functionality of distal IG/TR loci.

### Co-evolution of IG and TCR chains are associated with V gene counts

In total, the number of species with at least one fully assembled locus (both proximal and distal) varies from 36 to 43 across five types of immune loci (**Fig. 1A**). The number of species with two fully assembled and phased immune loci (both proximal and distal) varies from 18 to 21 for IGH, IGK, IGL, TRA loci and equals 33 for TRB loci (**Fig. 1A**). These numbers enable comparative analyses and suggest that the assembly complexity is comparable across all five types of adaptive immune loci. IGK loci were not detected in the five Chiroptera species, which is consistent with previous results (Pursell et al., 2024). Similarly, 2 out of 6 Rodentia species (the Merriam’s kangaroo rat, the Botta’s pocket gopher) did not have IGL loci, which is also consistent with previous studies describing short IGL loci in mice (Kos et al., 2022).

For all intact and proximal IG/TR loci, counts of productive V genes were compared. For Chiroptera species without IGK loci and Rodentia species without IGL loci, zero V genes were specified. **Fig. 1A** shows that the number of V genes per locus varies greatly across the species. The largest number of productive V genes was found in the brown big-eared bat IGH (640 V genes), the northern bat IGH (246 V genes), and the European snow vole IGH (217 V genes). All other loci contain less than 200 productive V genes. Counts of V genes positively correlate (all P-values < 0.01) with lengths of corresponding loci across all chain types (Fig. S2A).

Association analysis revealed links between V gene counts across some pairs of chains. First, counts of IGHV and IGLV genes positively correlate (here and further Pearson’s correlation is used, r=0.44, P=0.007; **Fig. 2A**). To assess whether the relationships between different chains and loci remains significant after accounting for shared evolutionary history, a phylogenetic linear model implemented in the *phylolm* package (Lam et al., 2014) with the lambda model (Freckleton et al., 2002) was used. Prior to the analysis, all values were log-transformed to ensure model fit. The detected correlation holds true after accounting for phylogenetic autocorrelation among the analyzed species (lambda=1, P=0.07). This result suggests that IGH and IGL loci co-evolve either in the response to one another, or in a shared genetic environment. While a similar relationship between IGH and IGK loci was not observed (Fig. S2B), the total IG light chain V gene (IGKV+IGLV) counts have a higher correlation with IGHV gene counts as compared to IGH/IGL (r=0.51, P=0.003, Fig. S2C). Counts of IGKV and IGLV genes show a stronger anticorrelation (r=–0.59, P=0.0003; **Fig. 2B**). This relationship was no longer significant when the slope was estimated with a phylogenetic regression (P=0.17), but the phylogenetic distribution of the counts appears discordant with the assumptions of phylogenetic regression (Uyeda et al., 2018) so we do not feel confident in the latter approach. We hypothesize that IGK and IGL chains play different roles across species, and expansion of one light chain locus leads to eventual contraction of another light chain locus. We also observed positive correlations between TRAV and TRBV gene counts (r=0.70, P=1.46྾10^−6^; **Fig. 2C**). This observation holds after accounting for phylogenetic autocorrelation (lambda=1, P=0.01). Because TRA and TRB loci encode α and β chains of the T-cell receptor, respectively, we assume that these loci co-evolve similarly to IGH/IGL loci.

**Figure 2.**
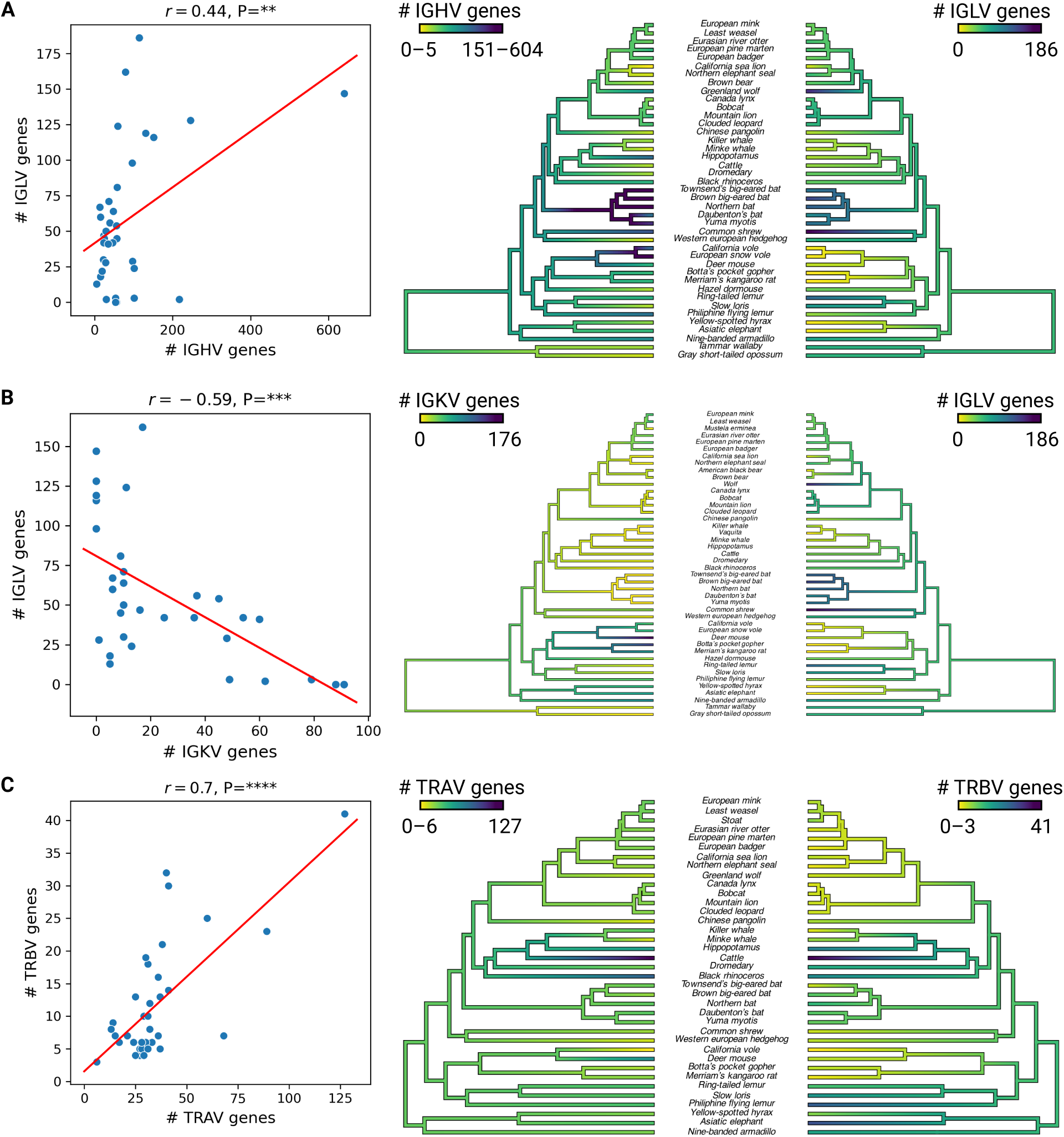
Correlations between productive V gene counts in proximal IG/TR loci. **(A)** Counts of IGHV and IGLV genes across 36 species. Only species with at least one reconstructed IGH locus and at least one reconstructed IGL locus were chosen. The Pearson’s correlation and the corresponding P-value are shown at the top of the plot. The red line shows the linear trend. The trees on the right show the species tree colored according to IGHV and IGLV gene counts, from yellow (0) to blue (the maximum V gene number). The species tree from Fig. 1 was used and the gene counts visualized with the contMap function from the phytools R package (Revell, 2024). **(B)** Counts of IGKV and IGLV genes across 32 species. **(C)** Counts of TRAV and TRBV genes across 37 species. Legends of (B) and (C) are consistent with (A).

### IG loci diverge faster than TR loci, and IGH loci diverge the fastest

To compare two sequences *S*_1_ and *S*_2_ representing loci of the same type of IG/TR loci, a new *locus distance* metric was introduced. First, local alignments between *S*_1_ and *S*_2_ using the YASS tool (Noé and Kucherov, 2005) were computed. Alignments longer than 5 kbp were analyzed, and the percentage of each of the sequences covered by these alignments was computed. The *locus similarity* of *S*_1_ and *S*_2_ was computed as the average of the corresponding fractions, and the *locus distance* was computed as 100 – the locus similarity. For each pair of loci of the same chain type, the distance between corresponding species was also calculated using the TimeTree5 database (Hedges et al., 2015). As expected, species distances positively correlate with locus distances for all IG and TR loci (r=0.81, P=<0.0001, **Fig. 3A**, Fig. S3); however, the locus distance values were distributed unevenly. Analysis of locus distances for the same species distances showed that IG loci were more diverged compared to TR loci (Kruskal-Wallis test, P=0.005); and IGH loci were the most diverged (Kruskal-Wallis test, P(IGH, IGK)=0.03, P(IGH, IGL)=1.22×10^−7^) (**Fig. 3B**). This suggests that IG loci evolve more rapidly and have fewer evolutionary constraints compared to TR loci. This might be explained by specifics of antigen recognition of antibodies and T-cell receptors: while antibodies bind antigens directly, TCRs also need to recognize the major histocompatibility complex representing antigen on the cell surface. Similarly, the faster divergence of IGH loci compared to IGK and IGL loci may reflect their distinct roles in antibody repertoires and respective evolutionary pressures. Previous studies report that IGH loci harbor more diverse V genes in primates (Yoo et al., 2024) and mice (Kos et al., 2022) and hypothesize that their primary role in antibody repertoires is to generate VDJ recombination diversity, while IGK and IGL chains protect against self-reactivity and maintain antibody stability (Collins and Watson, 2018).

**Figure 3.**
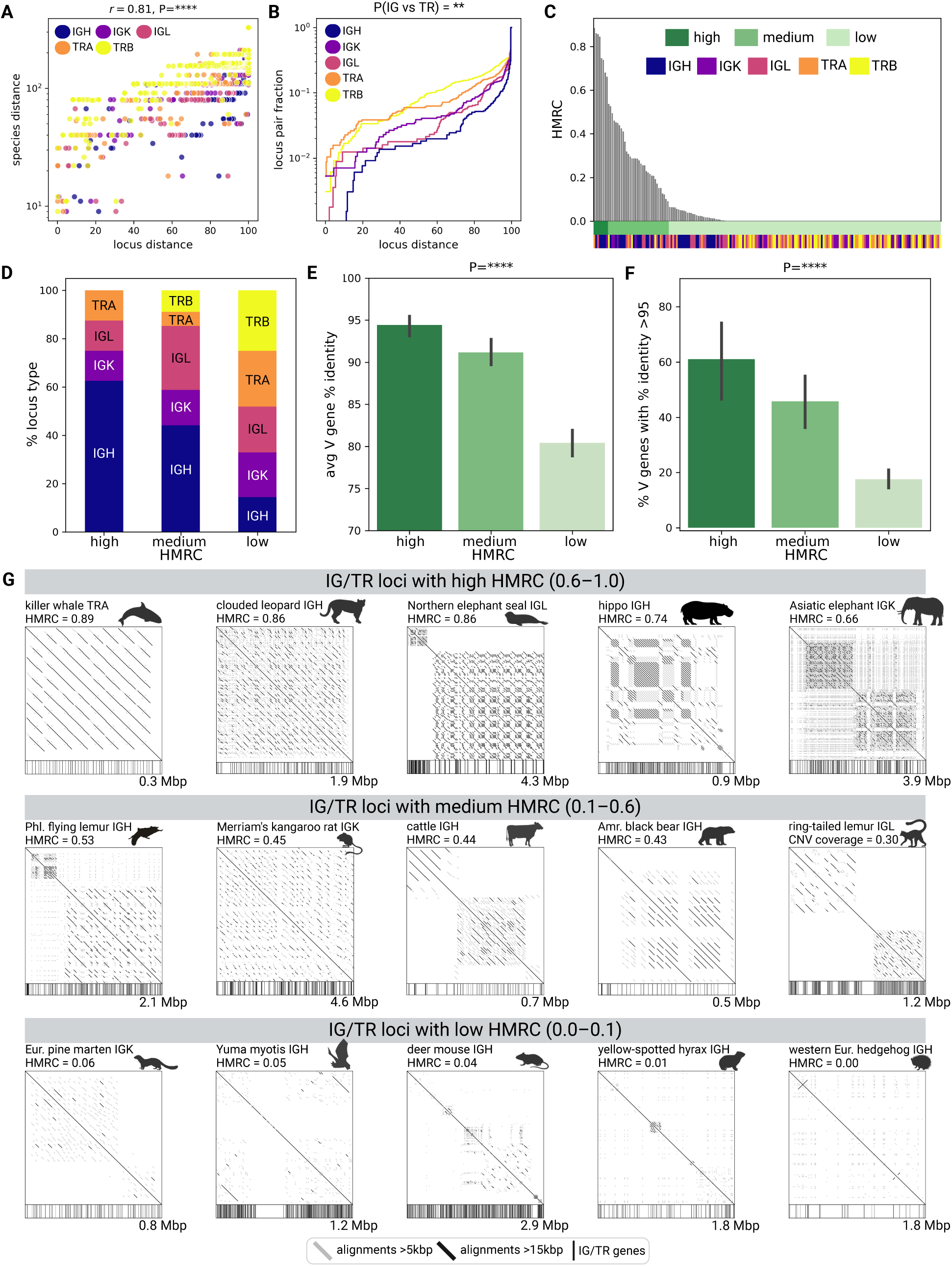
High multiplicity repeat content (HMRC) in IG/TR loci. **(A)** Locus distance vs species distance for all pairs of IG/TR loci of the same chain types. Points are colored according to the locus chain type: IGH (blue), IGK (purple), IGL (pink), TRA (orange), TRB (yellow). The species distances are shown in the logarithmic scale. Pearson’s correlation and P-value are shown at the top of the plot. **(B)** The cumulative histograms of locus distance values for five chain types. **(C)** HMRC of IG/TR loci sorted in descending order of the values. Each bar is colored in shades of green according to the HMRC class: high (dark), medium (medium), low (pale) and chain type (colors are consistent with **A**). **(D)** Fraction of each locus type in each of three HMRC classes. **(E)** Average percent identity of productive V genes in IG/TR loci across three HMRC classes. **(F)** The fraction of productive V genes with at least 95% similarity to another V gene in the same locus across three HMRC classes. **(G)** Examples of IG/TR loci with high, medium, and low HMRC. Each locus is shown as a dot plot, alignments longer than 15 (5) kbp are shown in black (gray). Positions of IG/TR genes are shown along the bottom of each plot.

### Some IG/TR loci are formed by high multiplicity repeats

Even though the direct comparison of IG/TR loci across species is often not possible, analysis of their genomic sequences allowed us to reveal their common features. Previous studies have shown that some IG/TR loci consist of large arrays of multiplicity repeats (e.g., the cattle IGH locus by Li et al., 2023), while others have local areas with repeat expansions (e.g., IGH loci of human and mouse, Sepulveda et al., 2005; IGH loci of great apes, Yoo et al., 2024) or are not repetitive at all (e.g., TRA loci of great apes, Yoo et al., 2024). To distinguish between these cases, the *high multiplicity repeat content (HMRC)* metric was developed and computed for each locus *S* as the fraction of positions in *S* covered by repeats of length at least *L* at least *N* times. Positions of repeats were computed as boundaries of the local alignments computed by YASS while aligning the locus versus itself. By default, *L*=15 kbp and *N*=5 (see “Computing high multiplicity repeat content of IG/TR loci” in Methods). The computed HMRC values were binned into three classes: high (HMRC: 0.6–1), medium (HMRC: 0.1–0.6), and low (HMRC: 0–0.1). Boundaries between classes were determined according to the largest relative drops of values in the sorted HMRC list. IG loci have higher HMRC compared to TR loci and IGH loci have the highest HMRC values (**Fig. 3C** & **Fig. 3D**). HMRC was not associated with V gene counts (Fig. S4A) nor locus lengths (Fig. S4B).

### IG/TR loci formed by high multiplicity repeats have a lower V gene diversity

To reveal features associated with repetitiveness, V gene similarities were analyzed. First, for each productive V gene, the closest V gene match within the same locus and the corresponding percent identity were computed. Then the average value of the best percent identities (*average percent identity*) was computed for each locus. The average percent identity is higher in IG compared to TR loci (Kruskal-Wallis test, P<0.0001; Fig. S5) and higher in IG/TR loci with high HMRC (95.1%) as compared to IG/TR loci with medium (92%) and low (80.2%) HMRC (Kruskal-Wallis test, P=3.06×10^−9^). This suggests that IG/TR loci with high HMRC harbor genes residing in copies of the repeat units (**Fig. 3E**). Moreover, IG/TR loci with high HMRC have a higher percentage of V genes with >95% similarity to another V gene within the same locus (average 61.0%) compared to IG/TR loci with medium (45.8%) and low (18.4%) HMRC (Kruskal-Wallis test, P=3.05×10^−9^, **Fig. 3F**). Both figures have a descending trend suggesting that coverage by high multiplicity repeats is an important factor contributing to V gene similarity. Visual analysis of examples of IG/TR loci with high HMRC shows that V genes reside in recent repeat units (**Fig. 3G**, the top line). In IG/TR loci with medium HMRC, units of repeats are either more diverged or cover a smaller fraction of the locus sequence thus resulting in V genes with lower similarity (**Fig. 3G**, the middle line). In IG/TR loci with low HMRC, IG genes do not belong to repeat copies and are potentially more likely to be dissimilar (**Fig. 3G**, the bottom line).

### IG/TR loci formed by high multiplicity repeats have a higher coverage by mobile elements

To reveal possible factors leading to high HMRC in IG/TR loci, the known classes of repeats were also computed using RepeatMasker (Chen et al., 2004). For each repeat type and each IG/TR locus, the percentage of the locus positions covered by repeats of a certain type was computed. The most abundant repeat classes are LINE/L1 (average locus coverage 18.8%), LTR/ERVL (1.9%), simple repeats (1.8%), LTR/ERVL-MalR (1.3%), LTR/ERV1 (0.9%), and low complexity sequences (0.5%) (**Fig. 4A**). Three types of know repeats revealed associations with HMRC: IG/TR loci with high HMRC have higher coverage by LINE/L1 (Kruskal-Wallis test, P=0.01) and LTR/ERV1 repeats (Kruskal-Wallis test, P=0.001) and lower coverage by low complexity regions (Kruskal-Wallis test, P=0.005) compared to IG/TR loci with low and medium HMRC (**Fig. 4B & 4C, Fig. S6**). These associations suggest that mobile elements might play a role in the formation of high IG/TR loci.

**Figure 4.**
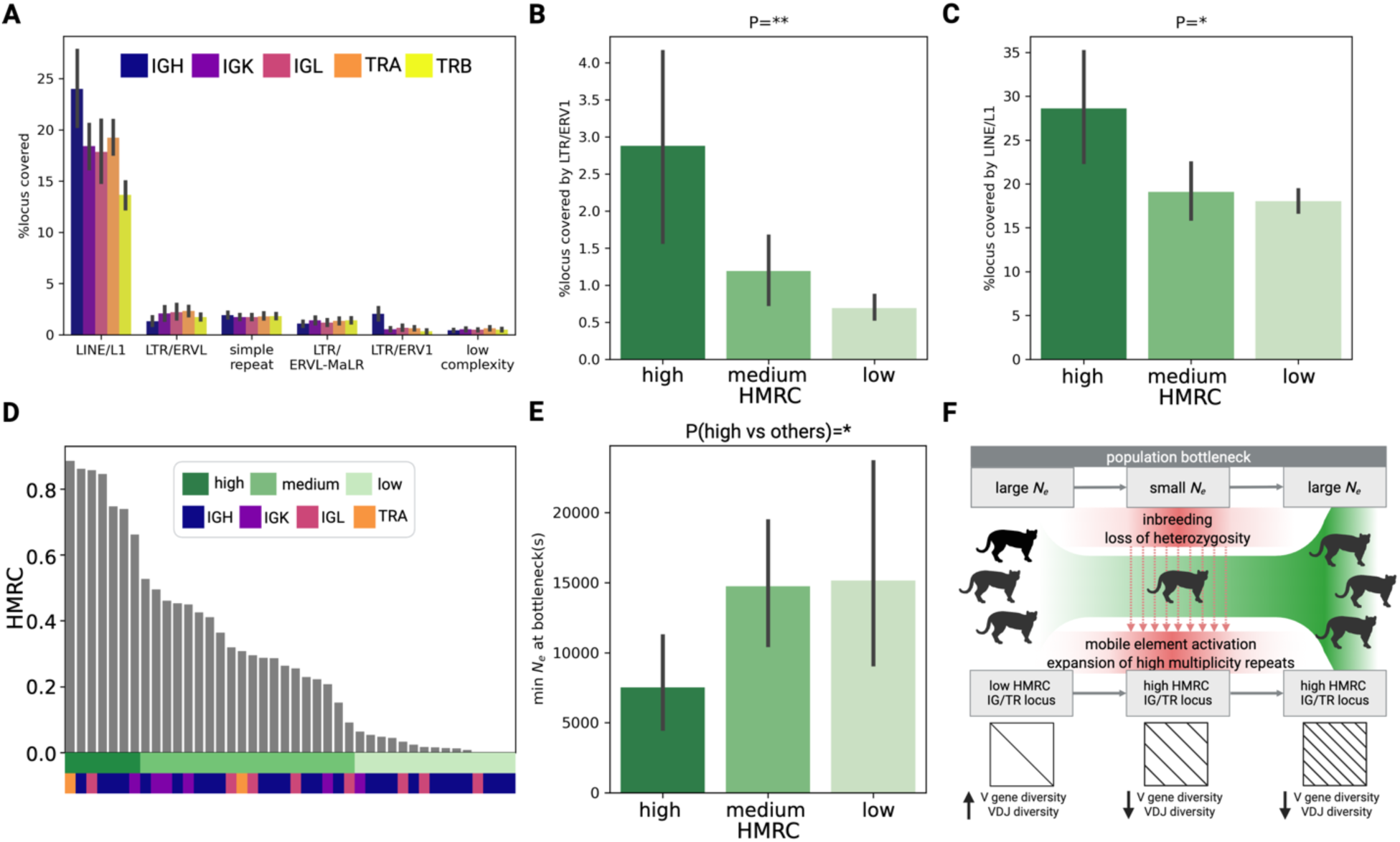
Characteristics of IG/TR loci with high, medium, and low HMRC. **(A)** Percentages of sequences covered by LINE/L1, LTR/ERVL, simple repeats, LTR/ERVL-MalR, LTR/ERV1, and low complexity repeats across five types of IG/TR loci. **(B)** Percentages of IG/TR loci covered by LTR/ERVL repeats across three HMRC classes. **(C)** Percentages of IG/TR loci covered by LINE/L1 repeats across three HMRC classes. **(D)** HMRC values (max HMRC value per species) sorted in descending order. Bars along the bottom show HMRC classes and the type of the corresponding locus. **(E)** The minimum *N_e_* value during population bottlenecks across three HMRC classes for non-closely related species. **(F)** A hypothesis showing links between population bottlenecks, loss of heterozygosity, activation of mobile elements, and high multiplicity repeat expansion in IG/TR loci.

### IG/TR loci with high HMRC are common for species that have undergone past population bottlenecks

Previous studies of plant genomes revealed elevated frequency of mobile element movements in inbred populations (Chen et al., 2020; De Kort et al., 2022). These studies suggest that inbreeding increases the fraction of recessive mutations and break mechanisms silencing mobile elements thus leading to structural diversification of the genome. To explore similar connections between IG/TR loci with high HMRC and population dynamics of the observed species, the effective population size (*N_e_*) was computed for each analyzed species using a modification of the PSMC model (Li & Durbin, 2011; see “Computing the effective population size” in Methods). For each species, *N_e_* values were computed over a period of time, and population bottlenecks were computed as local minimums of the *N_e_* curve and detected for 42 out of 44 species. We only considered past bottlenecks and used the *T_min_* threshold (by default, *T_min_*=50,000 years ago) for discarding bottlenecks occurring after it. The choice of *T_min_* is discussed in “Selecting a threshold for filtering out recent population bottlenecks” in Methods. For each species, the lowest *N_e_* value during a population bottleneck was identified. Because HMRC varies across loci with a species and previous studies suggested that structural diversification is a genome-wide effect, the maximum HMRC value and the corresponding HMRC class were selected across all five types of IG/TR loci for each species (**Fig. 4D**; **Table 2**). Species which IG/TR loci have high HMRC went through more severe bottlenecks with the average minimum *N_e_* 8598 as compared to species which IG/TR loci have medium and low HMRC with an average minimum *N_e_* of 14,123 (Fig. S7). To account for the impact of closely related species, two groups of species with similar loci were identified: IGH loci of felines (the clouded leopard, mountain lion, bobcat, and Canada lynx) and IGH loci of cetaceans (the vaquita and minke whale) (**Table 2**; Fig. S8). To minimize the impact of these groups, the average HMRC and the minimum *N_e_* at the bottleneck were computed for each of two groups; the HMRC class was assigned according to the binning in **Fig. 3C**. No pair of IG/TR loci had sufficient sequence similarity after collapsing these two groups. For the collapsed species, the observed pattern holds true with a P-value 0.04 (Mann-Whitney U test) (**Fig. 4E**). We suggest that mobile element movements increased during inbreeding have a pronounced influence on rapidly evolving IG and TR loci, leading to their structural diversification. While structural variations driven by mobile elements are often either neutral or deleterious at the genome-wide level (Solyom et al., 2012; Consuegra et al., 2021), this process may increase the genetic diversity of IG/TR loci through the generation of new gene-carrying repeat copies in populations suffering from losses of heterozygosity (**Fig. 4F**).

**Table 2.** Species with the maximum HMRC values across all five types of IG/TR loci. The corresponding locus and the selected HMRC value as well as the HMRC class are specified. Three groups of closely related species were shown as rows colored in green (cetaceans), yellow (felines), and blue (mustelids). For green and yellow groups, loci of the same type were selected. The minimum *N_e_* value of the brown bear was not computed because the genome assembly was reported as haploid.

| Sample ID | Common species name | Locus with the highest HMRC | HMRC | HMRC class |
| --- | --- | --- | --- | --- |
| mOrcOrc1 | Killer whale | TRA | 0.89 | high |
| mNeoNeb1 | Clouded leopard | IGH* | 0.86 | high |
| mMirAng1 | Northern elephant seal | IGL | 0.86 | high |
| mPumCon1.1 | Mountain lion | IGH* | 0.84 | high |
| mLynRuf1 | Bobcat | IGH* | 0.75 | high |
| mHipAmp2 | Hippopotamus | IGH | 0.74 | high |
| mEleMax1 | Asiatic elephant | IGK | 0.66 | high |
| mCynVol1 | Philippine flying lemur | IGH | 0.53 | medium |
| mManPen7 | Chinese pangolin | IGK | 0.50 | medium |
| mDasNov1 | Nine-banded armadillo | IGK | 0.46 | medium |
| mLynCan4 | Canada lynx | IGH* | 0.45 | medium |
| mDipMer1.0 | Merriam's kangaroo rat | IGK | 0.45 | medium |
| mBosTau | Cattle | IGH | 0.44 | medium |
| mUrsAme1.0 | American black bear | IGH | 0.43 | medium |
| mPhoSin1 | Vaquita | IGH* | 0.41 | medium |
| mBalAcu1 | Minke whale | IGH* | 0.36 | medium |
| mCanLor1 | Greenland wolf | IGL | 0.32 | medium |
| mChiNiv1 | European snow vole | TRA | 0.31 | medium |
| mLemCat1 | Ring-tailed lemur | IGL | 0.30 | medium |
| mMacEug1 | Tammar wallaby | IGH | 0.29 | medium |
| mCamDro1 | Dromedary | IGH | 0.29 | medium |
| mPleAur1 | Brown big-eared bat | IGH | 0.26 | medium |
| mSorAra2 | Common shrew | IGL | 0.26 | medium |
| mNycCou1 | Slow loris | IGH | 0.23 | medium |
| mDicBic1 | Black rhinoceros | IGH | 0.22 | medium |
| mThoBot1.0 | Botta's pocket gopher | IGK | 0.21 | medium |
| mEptNil1 | northern bat | IGH | 0.15 | medium |
| mCorTow1.0 | Townsend's big-eared bat | IGL | 0.09 | medium |
| mMarMar1 | European pine marten | IGK | 0.06 | low |
| mLutLut1 | Eurasian river otter | IGH | 0.05 | low |
| mMyoYum1.0 | Yuma myotis | IGH | 0.05 | low |
| mPerMan1.0 | Deer mouse | IGH | 0.04 | low |
| mMusNiv1 | Least weasel | IGL | 0.03 | low |
| mMyoDau2 | Daubenton's bat | IGH | 0.02 | low |
| mMonDom1 | Gray short-tailed opossum | IGL | 0.02 | low |
| mZalCal1 | California sea lion | IGH | 0.02 | low |
| mMusLut2 | European mink | IGH | 0.01 | low |
| mMelMel3 | European badger | IGH | 0.01 | low |
| mHetBru1 | Yellow-spotted hyrax | IGH | 0.01 | low |
| mMusErm1 | Stoat | IGL | 0.00 | low |
| mMusAve1 | Hazel dormouse | IGH | 0.00 | low |
| mMicCal1.0 | California vole | IGH | 0.00 | low |
| mEriEur2 | Western European hedgehog | IGH | 0.00 | low |

### IG loci accumulate more variations within a population compared to TR loci

To study the variation of IG/TR loci within a population even further, within each species and locus, we collected all pairs of homologous haplotype-resolved loci. Pairs of sequences where one locus is under assembled were discarded (see “Detecting under assembled haplotype-resolved IG/TR loci” in Methods). As a result, 95 haplotype-resolved pairs of IG/TR loci including 19 IGH, 15 IGK, 14 IGL, 20 TRA, and 27 TRB haplotypes were analyzed. To describe the level of variation between pairs of homologous haplotypes, *the haplotype similarity* metric was developed and employed. First, for each pair of haplotypes, local alignments were computed using YASS and sorted in descending order based on their lengths. For each haplotype in the pair, the sequence fraction covered by *n* longest alignments was computed for all *n* ranging from 1 to the total number of reported alignments (by default, *n* = 100). Computed fractions form a curve *F*(*n*), where *n* is the number of longest alignments. While the *F*(*n*) curves of similar haplotypes are expected to match (Fig. S9A), the *F*(*n*) curves of dissimilar haplotypes will have different shapes and areas under the curve (Fig. S9B). For each haplotype pair, the areas under two *F*(*n*) curves (*A*_1_ and *A*_2_) and the area between the curves (*A*_12_) were computed. Then *A*_12_ was subtracted from the average of *A*_1_ and *A*_2_, and the resulting value was referred to as the *haplotype similarity*. Because the maximum area under each curve is 100, haplotype similarity values vary from 0 (for diverged haplotypes) to 100 (for identical haplotypes). 95 haplotype pairs have haplotype similarity ranging from 21 to 100 (**Fig. 5A**), and TR haplotypes are more similar compared to IG haplotypes (Kruskal-Wallis test, P=2.7×10^−3^; **Fig. 5B**). IG haplotype similarity has a higher variance, which could be partially explained by the contribution of four taxonomic orders: the analyzed Artiodactyla and Carnivora species have more similar IG locus haplotypes compared to Chiroptera and Rodentia species (Kruskal-Wallis test, P=1.33×10^−6^; **Fig. 5C**). This pattern persists in TR pairs (Kruskal-Wallis test, P=7.81×10^−4^); however, haplotype similarity values are less distinct (Fig. S10). **Fig. 5D** showed that, within Chiroptera and Rodentia orders, diverged haplotypes can be completely dissimilar (e.g., the Townsend’s big-eared bat IGH) or characterized by dissimilar islands connected by more conserved fragments (e.g., the California vole IGH).

**Figure 5.**
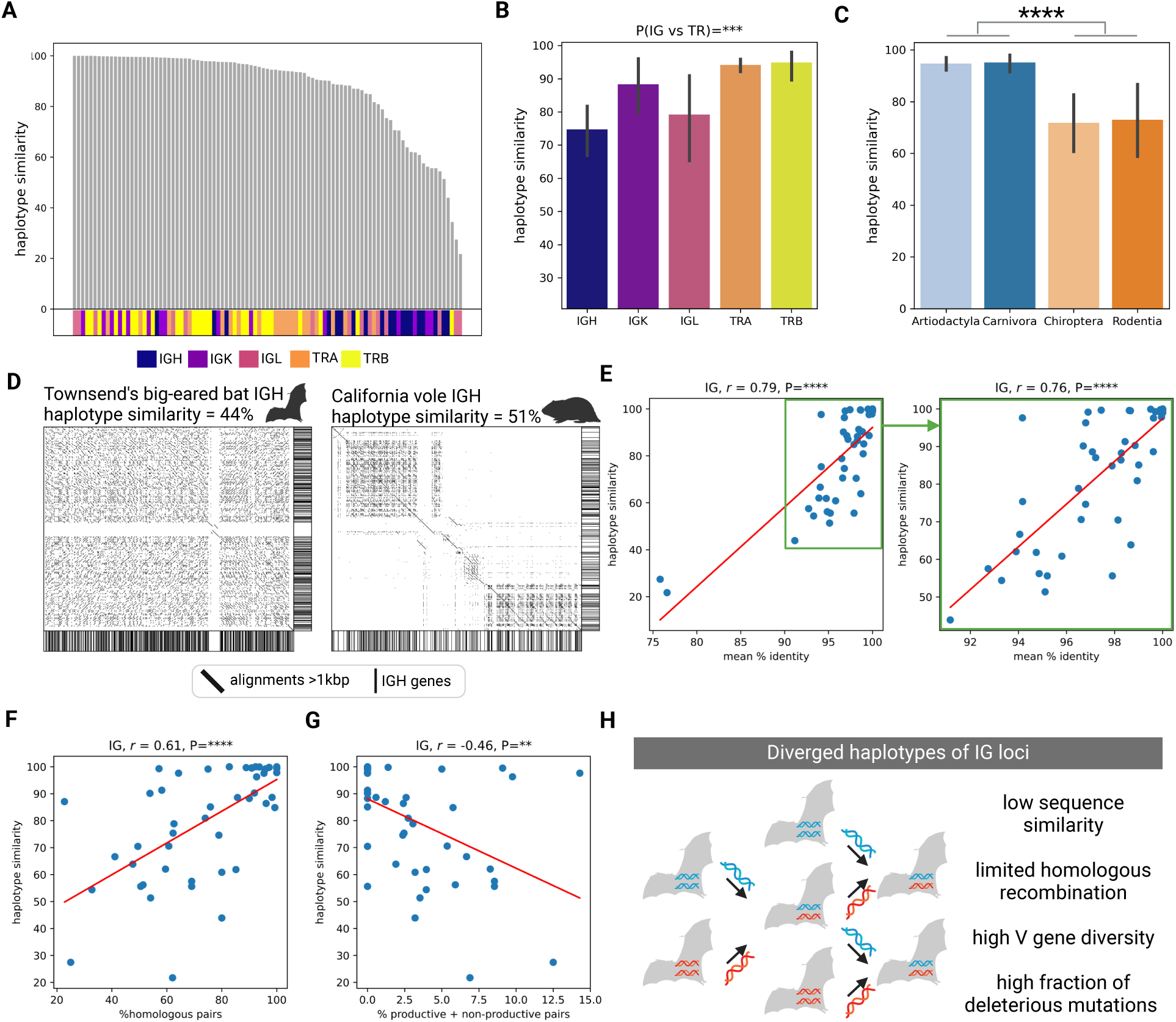
The variation of IG/TR locus haplotypes. **(A)** Haplotype similarities computed across 95 pairs of IG/TR haplotypes and sorted in descending order. The horizontal bar on the bottom shows IG/TR locus types. **(B)** Haplotype similarities across five types of IG/TR loci. P-value at the top shows differences between haplotype similarities values in IG and TR loci. **(C)** Haplotype similarities of IG haplotypes in four orders: Artiodactyla, Carnivora, Chiroptera, and Rodentia. P-value at the top corresponds to differences between combined Artiodactyla and Carnivora IG haplotype similarities vs combined Chiroptera and Rodentia IG haplotype similarities. **(D)** Dot plots of diverged haplotypes of Townsend’s big-eared bat IGH locus (left) and the California vole IGH (right). Only alignments longer than 1 kbp are shown. Bars along the bottom and the right of each plot shows positions of IGH genes. **(E)** Haplotype similarity vs average V gene similarity in IG haplotypes across all haplotype pairs (left) and all haplotypes without two outliers in the lower left corner (right). Pearson’s correlation and corresponding P-values are shown at the top of each plot here and further. **(F)** Haplotype similarity vs percentage of homologous V gene pairs across all IG haplotypes. **(G)** Haplotype similarity vs percentage of V gene pairs with one productive and one non-productive gene across all IG haplotypes. **(H)** A hypothetical scenario resulting in highly diverged IG haplotypes and their characteristics summarizing panels E-G.

### Diverged IG haplotypes might be subject to limited homologous recombination

For each productive V gene in a haplotype pair, the closest V gene in another haplotype and the corresponding percent identity of the alignment were computed. Only IG haplotypes were analyzed because of the higher variance of haplotype similarity values. For each haplotype-resolved IG locus, the average percent identity of V genes collected across both haplotypes was computed. The computed V gene similarities positively correlated with haplotype similarity values (r=0.79; P=1.59×10^−11^; **Fig. 5E**). **Fig. 5E** shows two outliers in the lower left corner (low gene similarity; low haplotype similarity), exclusion of which did not significantly change the correlation value (r=0.76, P=8.57×10^−10^). To reveal pairs of homologous V genes in each haplotype pair, the computed pairwise distances of V genes were combined into a distance matrix. The hierarchical clustering followed by flattening the dendrogram and extraction of clusters at the same cophenetic distance were applied to the matrix. The distance threshold was selected to maximize the number of clusters of size two. A cluster of size two was classified as homologous if it consisted of V genes from different haplotypes. The percentage of homologous pairs positively correlates with haplotype similarity (r=0.61, P=4.05×10^−6^; **Fig. 5F**). We also computed the percentage of clusters of size two where one V gene is productive and another is pseudogene and showed that it negatively correlates with haplotype similarity (r=–0.46, P=0.001; **Fig. 5G**). Correlations shown in **Fig. 5F**, 5G and the lack of homology between highly diverged IG haplotypes suggest that they might be excluded from chromosomal crossovers and inherited as one continuous allele (**Fig. 5H**). This possibly leads to an increase of V gene diversity, a decrease of homologous V gene pairs, an accumulation of V genes with broken copies. While mechanisms leading to the emergence of highly diverged IG haplotypes are unknown, we hypothesize that introgression (Zhang et al., 2016), suppressed recombination (Todesco et al., 2020), and / or elevated mutation rates (Xie et al., 2019) could play a role in their formation.

## Discussion

In this paper, five types of IG/TR loci (IGH, IGK IGL, TRA, and TRB) of 44 mammalian species across 13 taxonomic orders were characterized and compared. First, germline IG/TR genes were detected, and the gene counts across all pairs of IG/TR loci were compared. V gene counts in IGH and IGL loci showed a strong positive correlation suggesting that V genes of IGH and IGL loci may have evolved to generate preferential or more optimal pairs of heavy and light chain genes within expressed antibodies. A similar observation was made for TRA and TRB loci. These observations align with previous reports of preferential chain pairings in Abs and TCRs (DeKosky et al., 2016; Banach et al., 2021; Jaffe et al., 2022; Raybould et al., 2024) and might be important in future monoclonal antibody and TCR design studies.

On the contrary, V gene counts in IGK and IGL loci have a negative correlation, suggesting that the expansion of one locus leads to the contraction of another locus. These findings imply that immunoglobulin light chain loci serve different roles in B cell development and antigen recognition across mammalian species. For example, while human IGK locus is generally rearranged first, and IGL locus is rearranged if the IGK chain generates a non-productive or self-reactive rearrangement (Collins and Watson, 2018); IGL locus generates the majority of the initial antibody repertoire in pigs (Wertz et al., 2013). Follow-up studies of IGH, IGK, and IGL chain roles in mammalian antibodies will require expressed antibody repertoire sequencing data with paired chains (e.g., scRNA-Seq) and will be important for understanding the mechanisms of mitigating self-tolerance in antibodies and B-cell development across various species.

Pairwise analysis of IG/TR locus sequences shows that they evolve rapidly, which makes comparative analysis of their sequences through alignments difficult. Despite high sequence variability, some IG/TR loci share common organizational patterns. We observed that certain IG/TR loci consist of high multiplicity repeats, although the specific repeat units vary across species. Such IG/TR loci have a higher coverage by mobile elements and are more commonly found among species that went through severe population bottlenecks compared to non-repetitive IG/TR loci. This observation aligns with previous studies showing that inbreeding caused by population bottlenecks leads to activation of mobile elements and eventual structural diversification of the genome (Chen et al., 2020; De Kort et al., 2022) and suggests that a similar process takes place in inbred mammalian populations. Unlike other types of structural variations, high-multiplicity repeats can be detected using one locus sequence per species, and it is unclear whether other structural variations are associated with population bottlenecks. Generating population-wide genomics data across different species will be crucial for uncovering these evolutionary relationships.

IG/TR loci with high values of high multiplicity repeat content are characterized by a lower diversity of germline V genes because they reside in repeat copies. While a high diversity of germline genes is generally considered beneficial for boosting the diversity of naive adaptive immune repertoires (Bratsch et al., 2011; Schountz et al., 2017; Pennell et al., 2023), there is insufficient evidence to conclude that species with highly repetitive IG/TR loci have a reduced capacity to generate diverse immune responses. Further studies on expressed antibody and TCR repertoires are needed to address this question. Existing studies suggest that some species may have a higher diversity of D and J genes or evolved compensatory diversification mechanisms, such as ultralong cysteine-rich antibodies in bovines (Wang et al., 2013). Identifying similar mechanisms in other species could provide insight into the role of germline IG/TR diversity in shaping immune system versatility. If the germline diversity is shown to be a critical factor, it raises serious concerns as many species today are experiencing population declines due to human-induced habitat loss and fragmentation, potentially threatening the robustness of their immune systems.

Finally, analysis of haplotype-resolved IG/TR loci showed that some haplotypes do not share visible sequence similarity. Such loci are more prevalent in rodents and bats, and the mechanisms of their formation are unknown. Previously low sequence similarity of IG/TR loci and genes was reported for zoo animals (e.g., the Philippine flying lemur in Zhu et al., 2024) and model organisms (e.g., the mouse IGH, Watson et al., 2019; the macaque IGH, Peres et al., 2025) and suggests that human-mediated subpopulation mixing may be a contributing factor. Our analysis further showed that dissimilar haplotypes, while exhibiting high V gene diversity, also contain a higher proportion of broken V gene copies, suggesting that these loci may be partially or completely excluded from crossover events and thus accumulate pseudogenes at a faster rate. In this model, the entire IG/TR locus haplotype functions as a single continuous allele with a higher degree of linked polymorphisms. This observation raises important challenges for the development of IG/TR nomenclature in model organisms and opens new genomic questions. Specifically, it remains unclear how adaptive immune responses differ between species with highly diverged versus highly similar IG/TR haplotypes. Taking into account that conservation management strategies often involve subpopulation mixing, this question is particularly relevant to understanding how recovered populations adapt to wild environments.

In summary, IG/TR loci provide a unique perspective on genomic sequence diversity. Das et al., 2012 hypothesized that adaptive immune loci might have fewer evolutionary constraints due to V(D)J recombination. We expand on this idea, proposing that these regions are also highly susceptible to structural diversification, making them an ideal model for studying structural variations and answering important questions in fundamental and applied genomics. Further research on IG/TR loci and expressed adaptive immune repertoires both across and within populations will be essential for addressing these challenges and uncovering new insights into immune system evolution and function.

## Methods

### Modifying IgDetective and filtering broken IG/TR loci

The original version of IgDetective developed by Sirupurapu et al., 2022 was modified to detect both IG and TR V, D, J genes and determine boundaries of the respective loci. In addition to the RSS-based search described in the original work, we implemented an iterative procedure that aligns previously detected V genes to the input genome using minimap2 (Li., 2018) until no new V genes are detected. Groups of IG/TR genes separated by no more than 300 kbp were reported as IG/TR loci, and loci consisting of less than 10 V genes were discarded to filter out orphan genes. Boundaries of a locus were determined as 10 kbp before the leftmost IG/TR gene and 10 kbp after the rightmost position IG/TR gene. Loci with more than one contig per haplotype were discarded as under-assembled (Table S1). The only exception was made for bat species that have duplicated IGH loci (Pursell et al., 2024). Both copies of IGH loci were used for the analysis, and pairs of homologous IGH locus copies were found for the haplotype analysis.

### Computing high multiplicity repeat content of IG/TR loci

Because structural organization of the analyzed IG/TR loci is highly variable from species to species (and even within the same species), we computed a set of relaxed metrics *R*(*L*, *N*) showing the fractions of input sequence positions covered by repeats of length at least *L* at least *N* times. Such metrics allowed us to account for various types of segmental duplications and diverged repeat copies. *L*={10000, 15000} and *N*={3, 5} were tested and demonstrated high positive correlations with each other (Fig. S11). We thus used the strictest combination for the downstream analysis *R*(15000, 5). Further increasing *L* and *N* values resulted in zero values of *R*(*L*, *N*) for most loci and were discarded as non-informative.

### Computing the effective population size

The Pairwise Sequentially Markovian Coalescent (PSMC) model (Li and Durbin, 2011) was utilized to infer effective population size (*N_e_*) history for each mammalian genome assembly. The PSMC model is a widely adopted computational approach for demographic inference from a single genome. It analyzes the frequency of heterozygous sites throughout the genome to reconstruct the historical population size dynamics. Typically, heterozygous sites are identified by aligning short-read whole-genome sequencing data to a haploid genome assembly. However, modern Vertebrate Genomes Project (VGP) genome assembly pipelines (Rhie et al., 2021) primarily rely on PacBio HiFi sequencing, and Illumina short reads are not always produced. With haplotype-resolved genome assemblies, it is possible to directly identify all genetic differences between haplomes. To simplify the pipeline and leverage existing PSMC-compatible tools, we instead simulated perfect Illumina-like reads from the complete diploid genome assembly, employing a uniform coverage model with 10x coverage and a read length of 150 bp, these parameters were chosen to ensure compatibility and reliable performance of the existing tools. Subsequently, we followed the pipeline described by Li and Durbin, 2011: we aligned the simulated reads to either the primary haplotype or haplotype1, depending on the assembly curation, using BWA-MEM (Li, 2013). From these sorted alignments, we generated consensus sequences in FASTQ format using samtools (Danecek et al., 2021), capturing heterozygous positions between the two haplotypes through ambiguous nucleotide notation. The consensus FASTQ file was then processed using the internal PSMC utilities to generate the binary sequence input required for the PSMC analysis. Finally, we performed the PSMC analysis using the default parameters recommended by the developers. This pipeline efficiently enables the direct application of the PSMC model to diploid genome assemblies, eliminating the necessity for additional short-read sequencing data. To scale the PSMC trajectories, we applied parameters originally estimated for humans: a uniform mutation rate of 2.5×10^−8^ mutations per site per generation and a generation time of 25 years. These values were used consistently across all species in our study. We acknowledge that species-specific mutation rates and generation times would yield more accurate time scaling and absolute *N_e_* estimates. However, due to the lack of reliable estimates for many of the analyzed species, we opted to use uniform parameters to facilitate consistent comparative purposes.

### Selecting a threshold for filtering out recent population bottlenecks

The Ne curves were computed in the time frame from 0 to 27M years ago. To identify the minimum *T_min_* threshold, we tested values starting from 10,000 to 200,000. For all *T_min_* values, species with IG/TR loci with the high HMRC have lower minimum Ne at bottlenecks compared to the medium and high classes (Fig. S12). We thus selected the first value (*T_min_* = 50,000) resulting in a statistically significant association.

### Detecting under assembled haplotype-resolved IG/TR loci

Visual assessment of dot plots computed for IG/TR haplotypes revealed that some pairs represent potentially broken assemblies even if each haplotype has only one contig for the corresponding IG/TR locus. In most cases, we were able to make this conclusion because a noticeable part of the prefix or suffix of one of the contigs in the pair was not aligned to another contig in the pair (Fig. S13). Because the described haplotype similarity metric will be artificially low in such cases, we excluded such pairs from the analysis. In total, 20 out of 115 haplotype pairs were excluded from the analysis using this procedure.

## Contributions

MPo, CWT, KPK, MPe, and YS conceptualized the study. MPo, KV, AZ, AB, MPe, and YS designed experiments and performed computational analyses. MPo and YS wrote the manuscript. All authors have approved the text and provided comments to it.

## Acknowledgments

KV and MPe are supported by National Institute of General Medical Sciences (R35GM151348).

## Declaration of generative AI and AI-assisted technologies in the writing process

During the preparation of this work the author(s) used Chat GPT in order to improve language and readability. After using this tool/service, the author(s) reviewed and edited the content as needed and take(s) full responsibility for the content of the publication.

## Data availability

The new version of IgDetective is publicly available at GitHib: github.com/Immunotools/IgDetective. Scripts and computed characteristics of species and loci (*N_e_* statistics, locus lengths, gene counts, similarity metrics) are available at github.com/posma/ComparativeImmunogenomics.

