## Supplemental Materials for "Comparative analysis of mammalian adaptive immune loci revealed spectacular divergence and common genetic patterns"

### Supplemental Figures

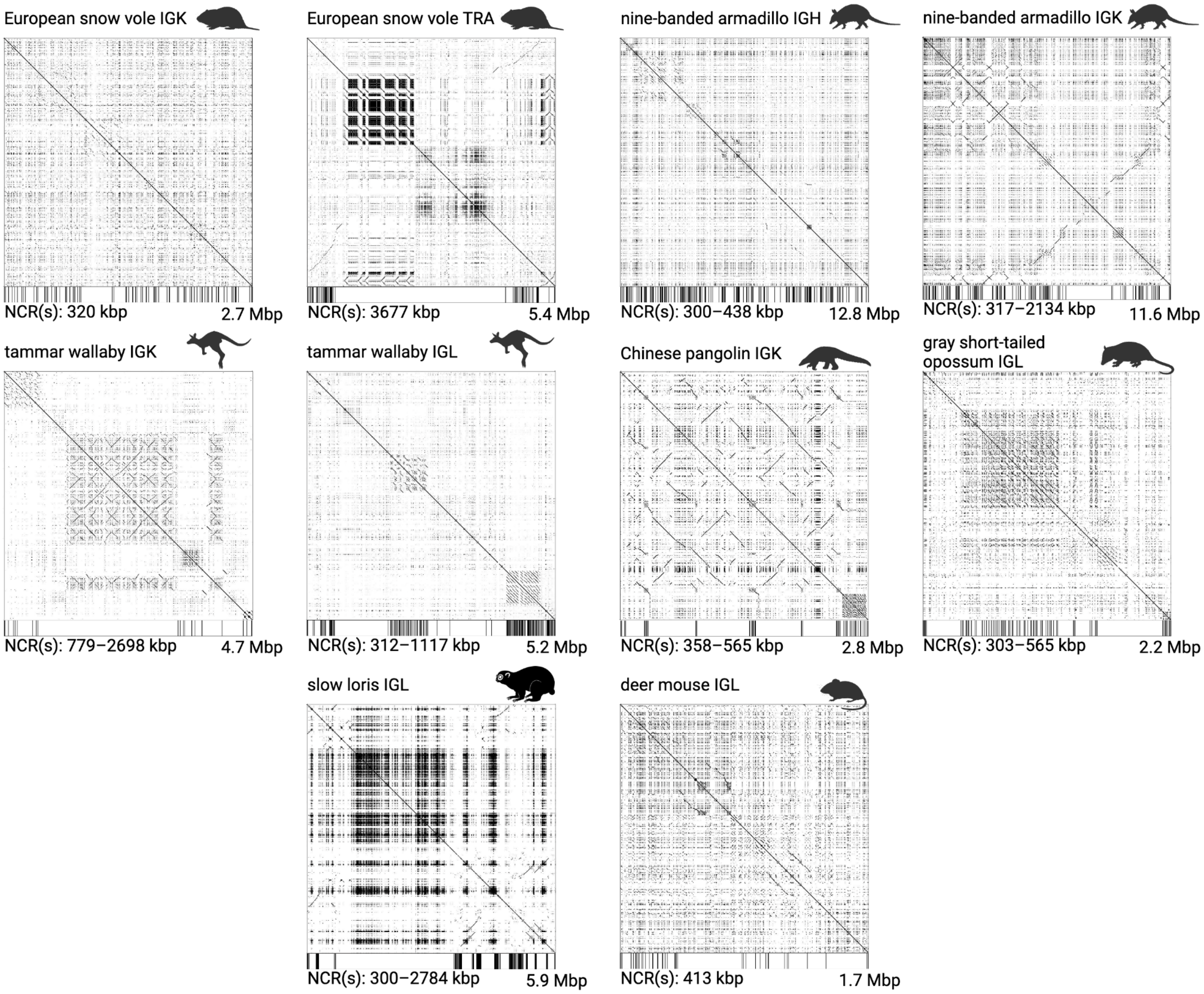


**Figure S1. The dot plots of ten distal IG/TR loci.** Positions of V genes are shown as a horizontal bar on the bottom of each plot. NCRs (non-coding regions) refer to regions between the proximal parts. The total locus lengths and NCR length ranges are specified on the bottom of each plot.

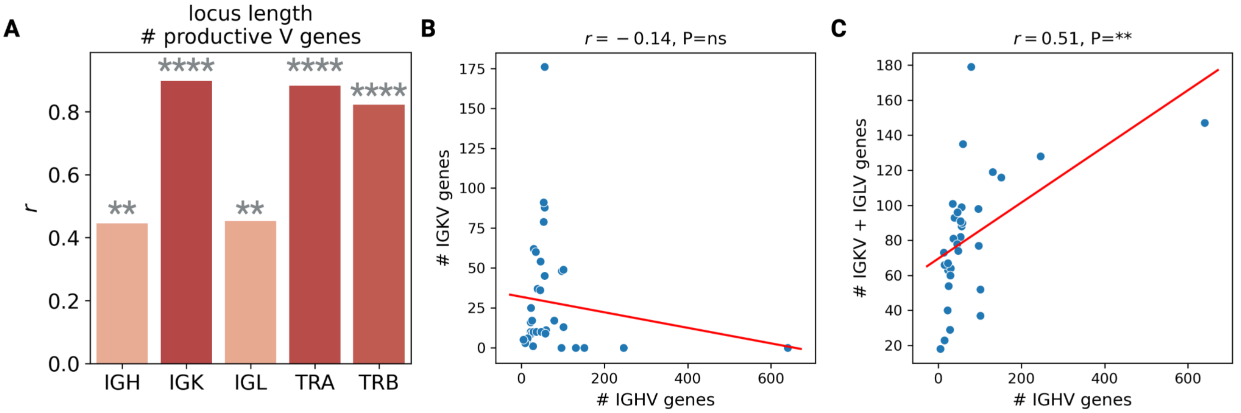


**Figure S2. Correlations between productive V counts and IG/TR locus lengths. (A)** Pearson's correlations between locus lengths and V gene counts across five types of IG/TR loci. Significance levels of correlation P-values are shown at the top. **(B)** Counts of IGHV and IGKV genes across 36 species. **(C)** Counts of IGHV and IGKV + IGLV genes across 32 species. Legends of (B) and (C) are consistent with **Fig. 2A**.

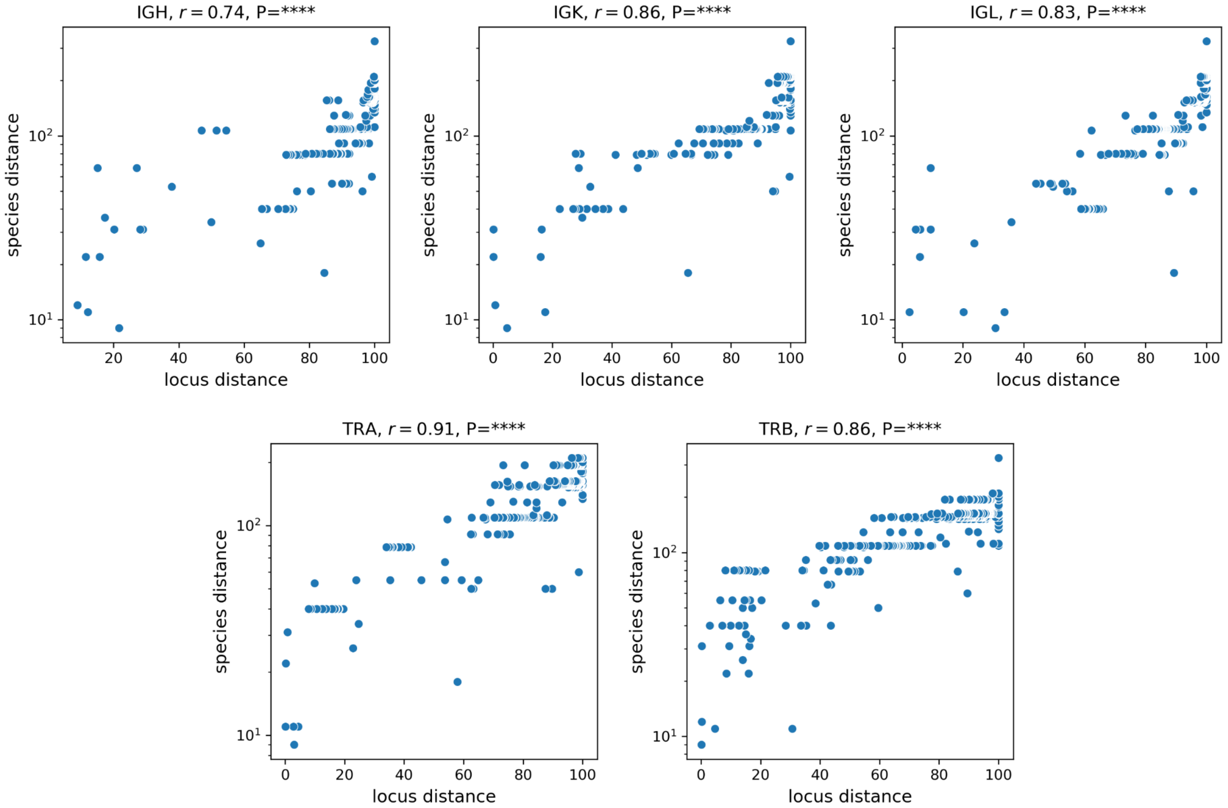


**Figure S3. Distances between pairs of species and respective pairs of IG/TR across all five chain types.** The *r* value at the top of each plot corresponds to Pearson's correlation value. P-values are $2.91\times10^{-160}$ (IGH), $3.12\times10^{-185}$ (IGK), $5.79\times10^{-199}$ (IGL), $1.32\times10^{-263}$ (TRA), $2.86\times10^{-248}$ (TRB).

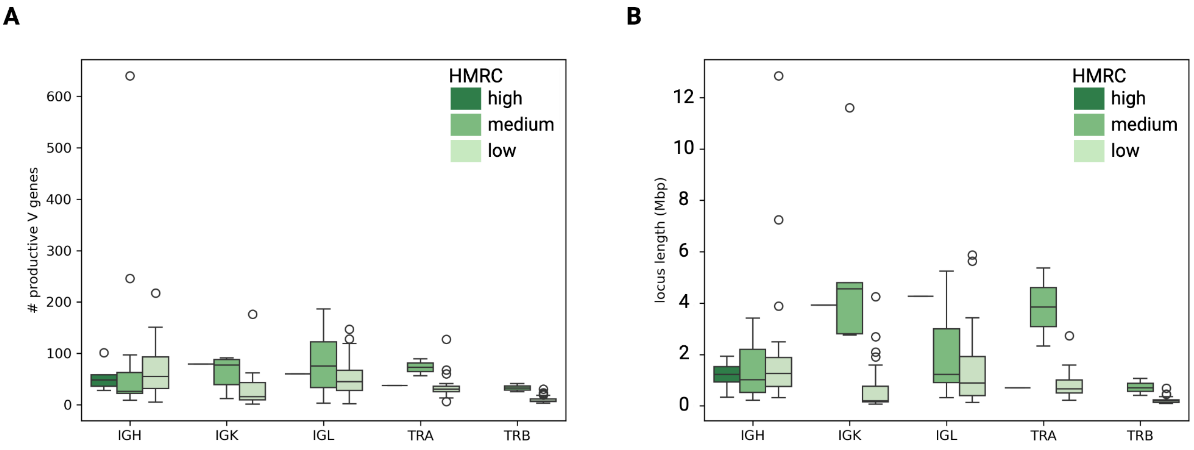


**Figure S4. HMRC classes vs counts of productive V genes (A) and locus lengths (B) across all types of IG/TR loci.**

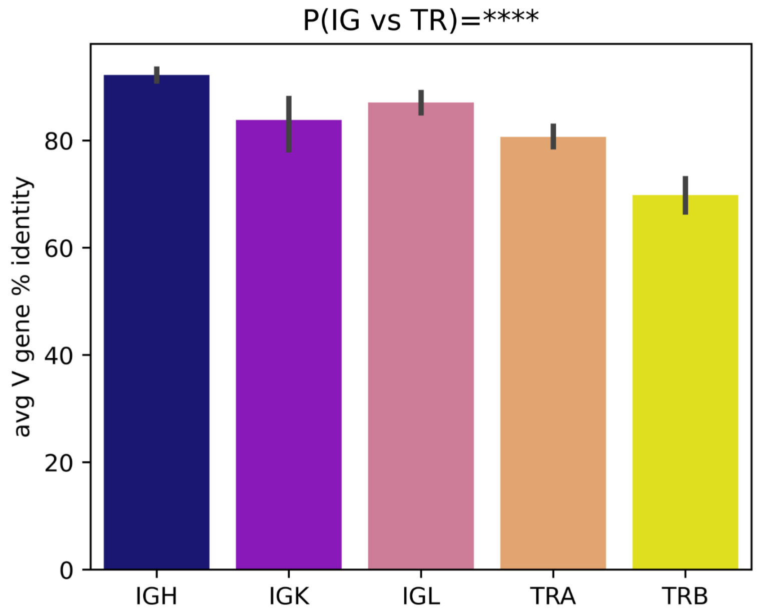


**Figure S5. Average percent identity of productive V genes across IG/TR loci.**

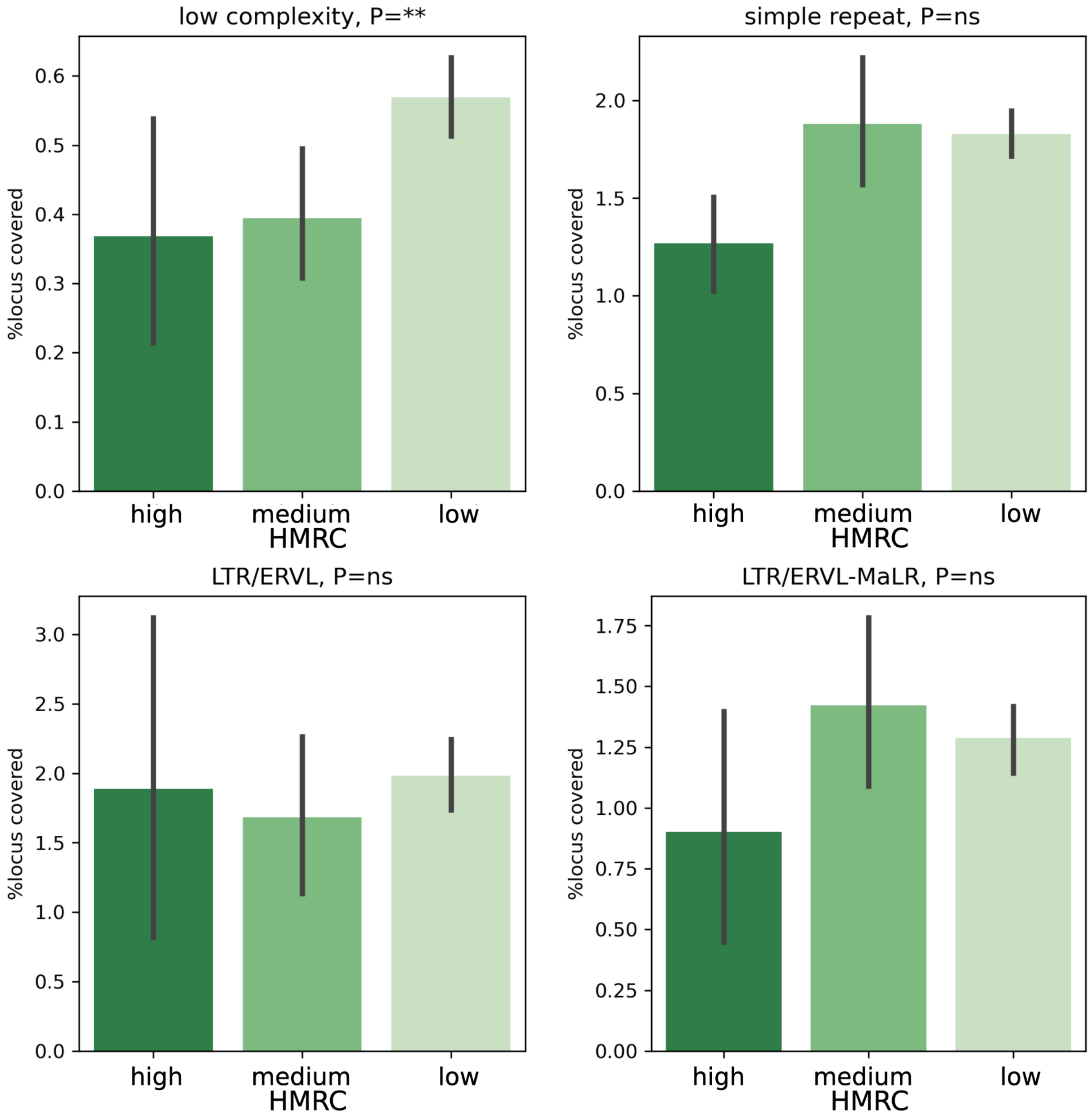


**Figure S6. Percentages of locus positions covered by low complexity repeats, simple repeats, and repeat classes LTR/ERVL, LTR/ERVL-MaLR vs HMRC classes.**

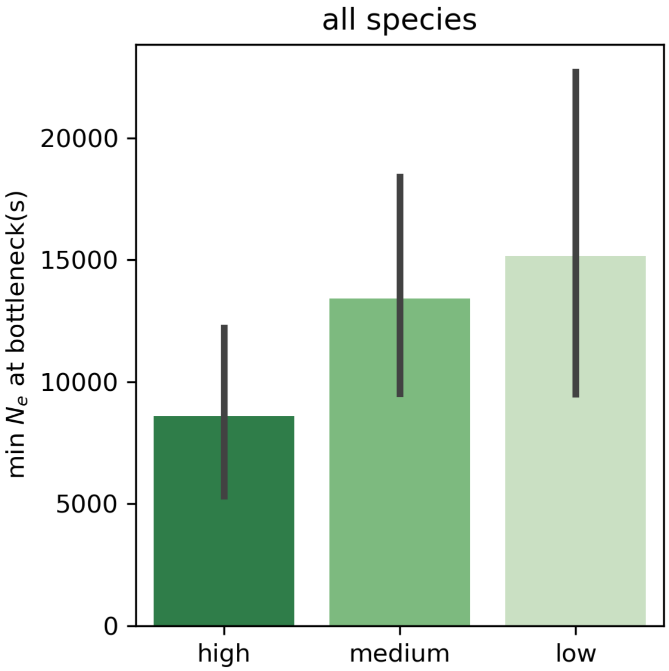


**Figure S7. The minimum $N_e$ value at population bottlenecks across three HMRC classes for all species.**

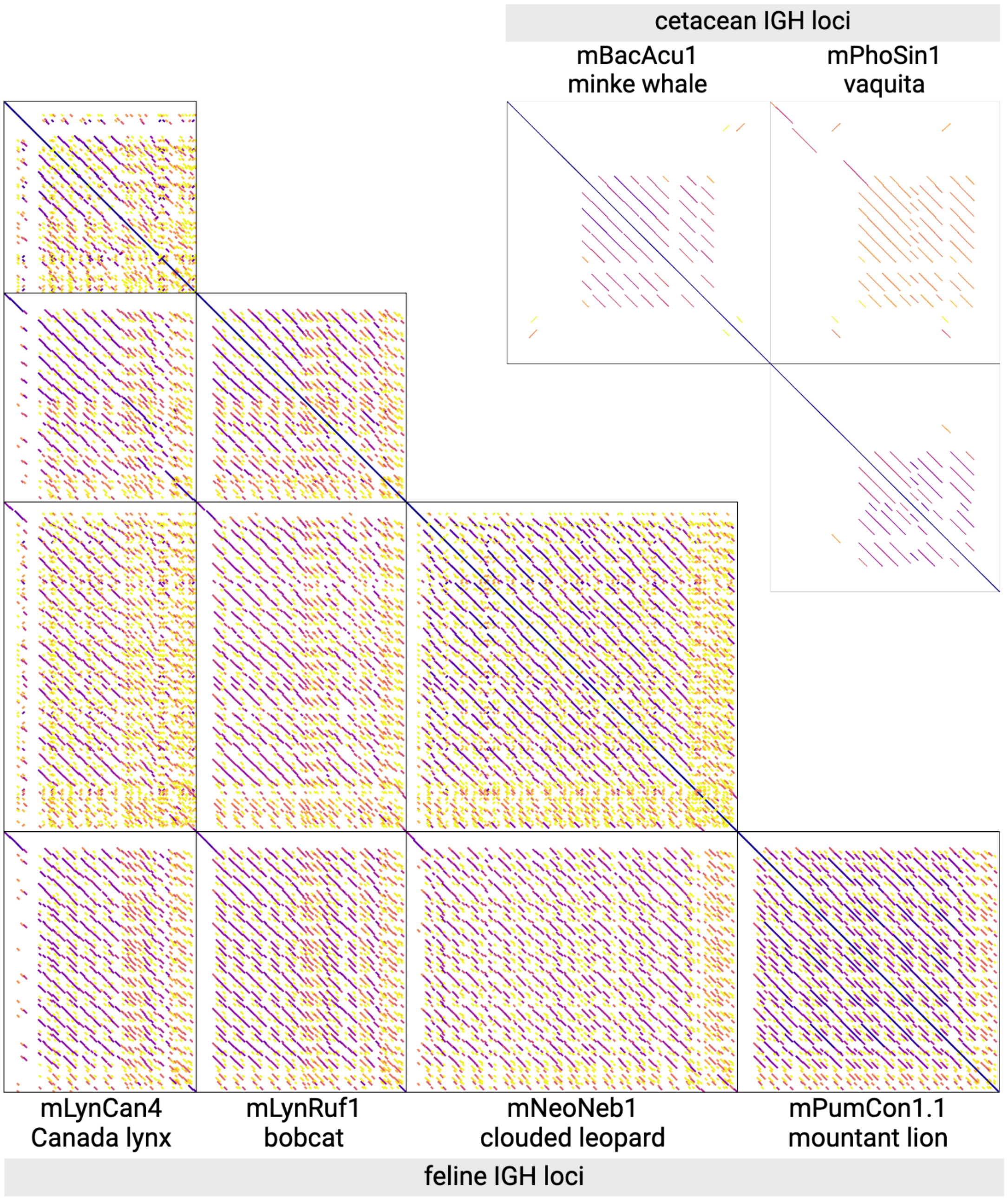


**Figure S8. Pairwise alignments of IGH loci of two groups of closely related species: felines (the lower triangle) and cetaceans (the upper triangle).** Alignments are colored according to their percent identities: from yellow (≤85%) to dark violet (100%). Dot plots were generated using the PatchWorkPlot tool (Pospelova and Safonova, 2025).

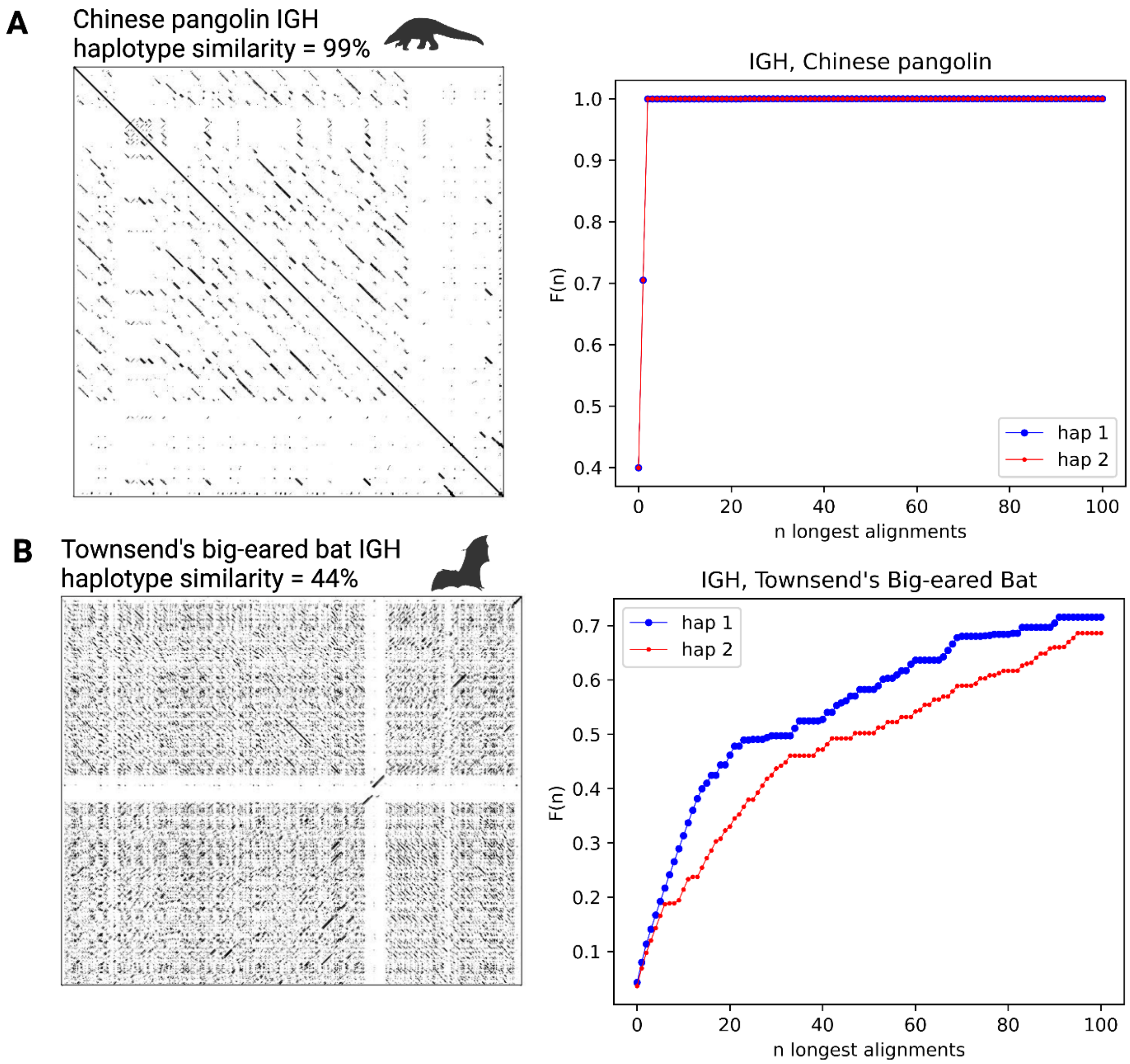


**Figure S9.** ***F(n)* curves and corresponding haplotype similarity values. (A)** Similar haplotypes of the Chinese pangolin IGH. **(B)** Diverged haplotypes of the Townsend's big-eared bat IGH.

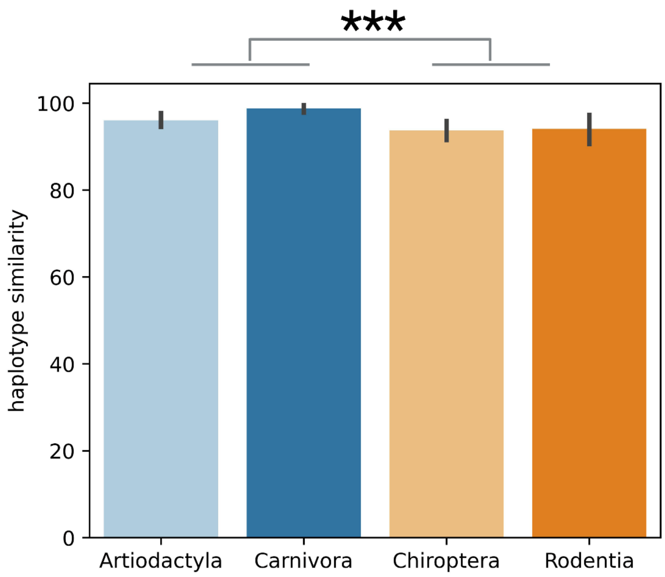


**Figure S10. Haplotype similarities of TR loci in Artiodactyla and Carnivora species vs Chiroptera and Rodentia species.**

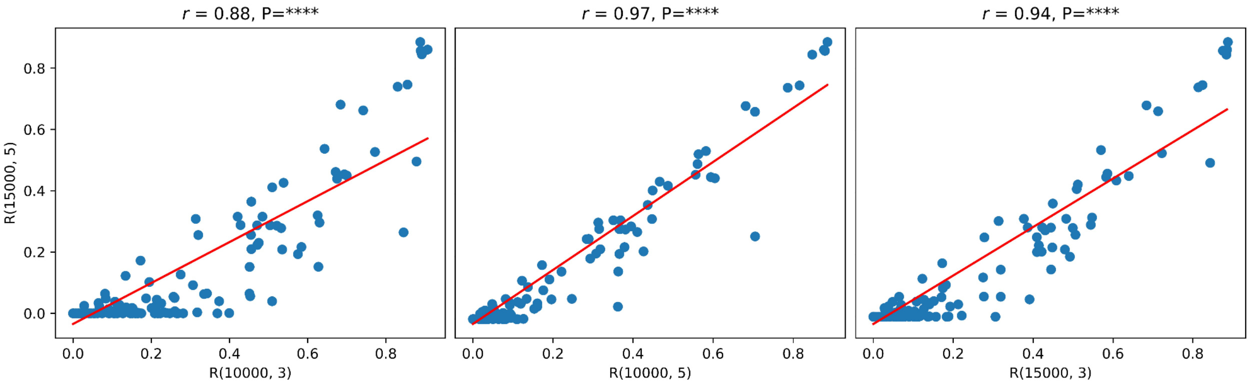


**Figure S11. Comparison of R(10000, 3), R(10000, 5), R(15000, 3) values with R(15000, 5) values.** *R* and P values at the top of each plot show Pearson's correlation and corresponding P-values.

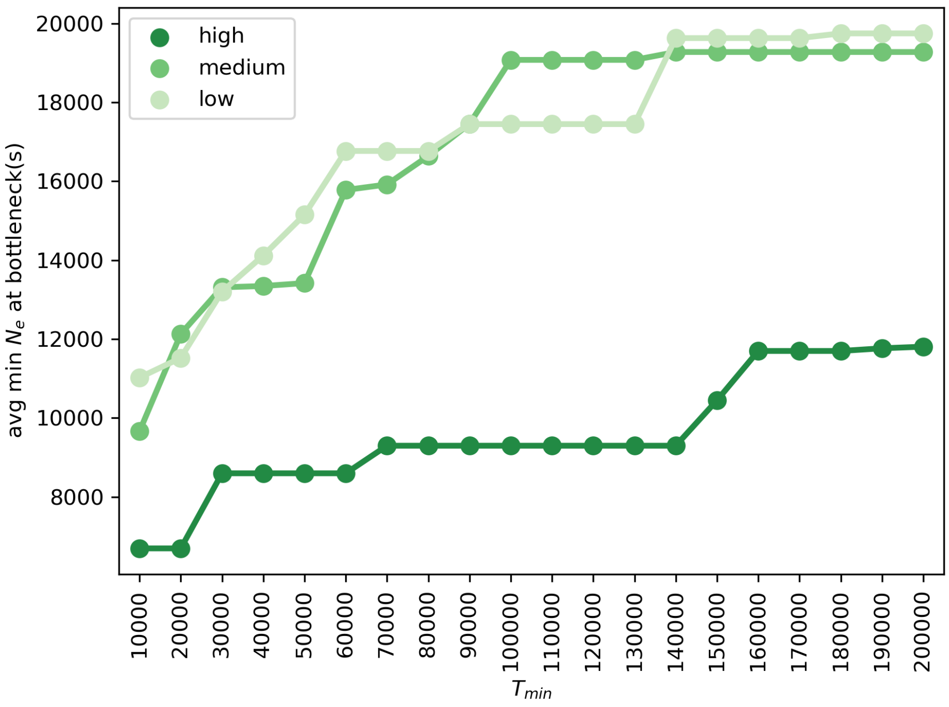


**Figure S12. The minimum average $N_e$ value at population bottlenecks across three HMRC classes for $T_{min}$ values ranging from 10,000 to 200,000 years ago.**

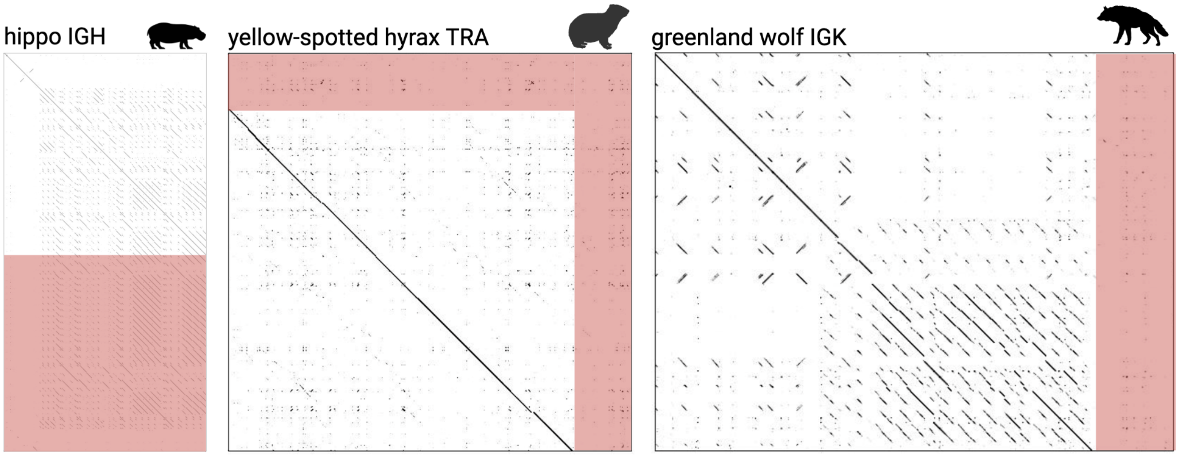


**Figure S13. Examples of under assembled haplotype pairs.** Red areas correspond to sequence fragments with no alignment and no assembly in another haplotype.
